# A Neuroeconomic Framework for Creative Cognition

**DOI:** 10.1101/184754

**Authors:** Hause Lin, Oshin Vartanian

## Abstract

Neuroeconomics is the study of the neurobiological bases of subjective preferences and choices. We present a novel framework that synthesizes findings from the literatures on neuroeconomics and creativity to provide a neurobiological description of creative cognition. It proposes that value-based decision-making processes and activity in the locus coeruleus-norepinephrine (LC-NE) neuromodulatory system underlie creative cognition, as well as the large-scale brain network dynamics shown to be associated with creativity. This framework allows us to re-conceptualize creative cognition as driven by value-based decision making, in the process providing several falsifiable hypotheses that can further our understanding of creativity, decision making, and brain network dynamics.

---

According to the standard definition, products that are both novel and useful within a given context are considered creative (Diedrich, Benedek, Jauk, & Neubauer, 2015; Runco & Jaeger, 2012; see also Sternberg, 1999). However, despite notable recent advances in the neuroscience of creativity (for reviews see Jung & Vartanian, 2018; Vartanian, Bristol, & Kaufman, 2013) and a wealth of correlational data from brain imaging studies (for meta-analyses see Boccia et al., 2015; Gonen-Yaacovi et al., 2013; Wu et al., 2015), a critical question that remains unanswered is how the brain produces ideas that satisfy the above two criteria. Part of this shortcoming may be due to the lack of mechanistic accounts of brain processes that underlie creative cognition.

We work from the assumption that a complete account of creativity will require not only an understanding of its cognitive architecture, but also the neural systems that underlie it. Towards that end, we propose a novel and neurologically plausible framework for creative cognition. This framework has three components: First, it proposes a neuroeconomic approach to creativity, suggesting that value-based decision-making processes underlie creative cognition. Second, it describes how the locus coeruleus-norepinephrine (LC-NE) neuromodulatory system could support creative cognition by adaptively optimizing long-term subjective value or payoff associated with preferences and/or choices. Third, it suggests that the dynamic interactions within and between brain networks observed during creative cognition are driven by activity in the LC-NE system and the interconnected brain regions that compute and evaluate subjective value. By bringing together a diverse range of findings from different fields, this framework provides a new conceptualization of creative cognition as driven by value-based decision making. It also points the way to future research by providing novel and testable hypotheses that are relevant to the fields of creativity, decision making, and brain network dynamics.

## Value-based decision-making processes underlie creative cognition

### Neuroeconomics of creative cognition

Neuroeconomics is a young but thriving interdisciplinary field that investigates the neurobiological processes underlying subjective preferences and choices (Camerer, 2013; Rangel, Camerer, & Montague, 2008; Konovalov & Krajbich, 2016). Specifically, it focuses on the computations the brain carries out to make value-based decisions, as well as the biophysical implementation of those computations (Bogacz, Brown, Moehlis, Holmes, & Cohen, 2006; Tajima, Drugowitsch, & Pouget, 2006; Wang, 2002). Value-based choices are pervasive in everyday life, ranging from the mundane to the consequential. Essentially, any choice that requires us to express our subjective preferences and to choose from among two or more alternatives is a value-based choice (e.g., Do you want an apple or an orange? Do you prefer the universe or the multiverse model?). These choices often lack an intrinsically correct answer, and depend instead on subjective preferences. They are called *valued-based* or *economic* choices because most neurobiological models of decision making have integrated economic constructs such as value maximization into their frameworks. These models assume that decision makers make choices by assigning a value to the available options, and then select the option with the highest value (Kable & Glimcher, 2009; Padoa-Schioppa, 2011; Rangel et al., 2008).

The basic premise of the present framework is that creative cognition is similarly supported by value-based decision-making processes. In its stronger form, creative cognition would be considered just another form of value-based decision making, because it too is underwritten by the same neural systems that drive value computations in the context of making choices about other commodities (e.g., material goods). That is, process-wise, creative cognition resembles making choices in everyday settings because it too involves generating multiple ideas and then selecting the idea with the highest subjective value amongst generated ideas (see Vartanian, 2011). We use the term subjective value in its traditional economic sense (i.e., the total amount of satisfaction that a good or service brings about) rather than how it is sometimes used within the creativity literature (i.e., to imply the usefulness of an idea, see Harrington, 2018). The notion of subjective value is central to choice theories in many disciplines including ecology, economics, and psychology, serving as an integrated decision variable by which options are compared (Padoa-Schioppa, 2011; Pearson, Watson, & Platt, 2014; Rangel et al., 2008).

The value of a creative idea or product therefore refers to the overall satisfaction derived from that idea or product, and is critical for driving choice behavior. Creative ideas will be assigned higher values and will be more likely to be selected if they maximize overall satisfaction, depending on how highly they score on attributes such as (but perhaps not limited to) novelty and usefulness within a given context. In this sense, the underlying process is similar to what might occur in other decision contexts like dietary choice, where a food will be assigned higher value if it were to score high on attributes such as healthiness and taste (e.g., Hare, Camerer, & Rangel, 2009). Consistent with the ideas of philosopher Paul Souriau (as cited in Campbell, 1960, p. 386) who noted that “of all of the ideas which present themselves to our mind, we note only those which have some *value* and can be utilized in reasoning” (italics added), the basic premise of our model is that a domain-general machinery that computes value is central to making choices in many contexts, including those that require creative thinking.

Importantly, our framework is agnostic as to which specific attributes should be used to evaluate the value associated with creativity (e.g., novelty, usefulness, surprise, etc.). Value in any context is simply determined by the following formula: value = Σ weight*attribute + error (Berkman et al., 2017). Depending on the researcher’s theory and/or decision context, different combinations of attributes and associated weights could be entered into the equation. In addition, our framework does not equate creativity with value. Rather, value is a way to conceptualize and think about how people *judge* or *evaluate* creative ideas, products, or solutions. As with any other decision studied by economists, we are proposing that value is an assessment or evaluation of a good or product—in this case an idea, product, or solution. Finally, the combination of attributes and weight used to compute value could take additive or multiplicative forms. In this sense the framework offers the necessary flexibility for representing and testing different theoretical approaches using a common (i.e., value-based) metric. Accordingly, depending on the attributes, weights and combinatorial setup entered into the equation, one will in turn observe different patterns of neural activity.

Within the psychological literature, the idea that creativity involves thought processes that resemble value-based decision making is not without precedent. One well-known example is the family of Blind Variation Selective Retention models, in which creativity includes generation and selection steps, the latter of which explicitly incorporates evaluative processes (Basadur, Graen, & Green, 1982; Campbell, 1960; Simonton, 1999; see also Vartanian, 2011). Specifically, after an initial step that involves the generation of candidate ideas, the second step involves the engagement of evaluative process for selecting the best idea(s) (based on certain criteria) for further consideration. Importantly, the term “blind” simply indicates insufficient prior knowledge about an idea’s usefulness (imonton, 2016). Another example is Sternberg and Lubart’s investment theory of creativity (Lubart & Sternberg, 1995; Sternberg, 2006, 2012), according to which creative people excel at pursuing and further developing ideas that have growth potential, but happen to be unknown or out of favor within the field. In this sense, they “buy low and sell high in the realm of ideas” (Sternberg, 2012, p. 5). Creative idea generation therefore involves evaluative processes that help select unpopular ideas for further nurturing. However, although both Blind Variation Selective Retention models and the investment theory of creativity acknowledge a relationship between value maximization and creative cognition, they do not provide neurobiological and mechanistic descriptions of how value maximization contributes to creativity. In what follows we will review evidence suggesting a relationship between value-based decision making and creativity, and will argue that the former helps to realize the latter.

One of the most robust findings from neuroeconomic research is that across species and studies, a specific set of brain regions, including the ventromedial prefrontal cortex (vmPFC), the orbitofrontal cortex (OFC), posterior cingulate cortex (PCC), and the striatum, are involved in value-based decision making (Padoa-Schioppa & Cai, 2011; Padoa-Schioppa & Conen, 2017; Rangel et al., 2008; Rich & Wallis, 2016) (Figure 1). For example, functional magnetic resonance imaging (fMRI) studies have shown that blood-oxygen-level dependent (BOLD) signals in the vmPFC correlate with behavioral preferences for beverages (McClure et al., 2004) and the subjective value of delayed monetary rewards (Kable & Glimcher, 2007; McClure, Laibson, Loewenstein, & Cohen, 2004). Crucially, converging evidence from fMRI (Bartra, McGuire, & Kable, 2013; Clithero & Rangel, 2014; Grueschow, Polania, Hare, & Ruff, 2015), lesion (Buckley et al., 2009; Camille, Griffiths, Vo, Fellows, & Kable, 2011; Hogeveen, Hauner, Chau, Krueger, & Grafman, 2016), and electrophysiological (Padoa-Schioppa, 2011; Padoa-Schioppa & Assad, 2006; Rich & Wallis, 2016) studies suggests that a set of brain regions comprised of the OFC, vmPFC, medial prefrontal cortex (mPFC), and PCC not only represents value, but also evaluates choice alternatives during value-based decision making.

**Figure 1.**
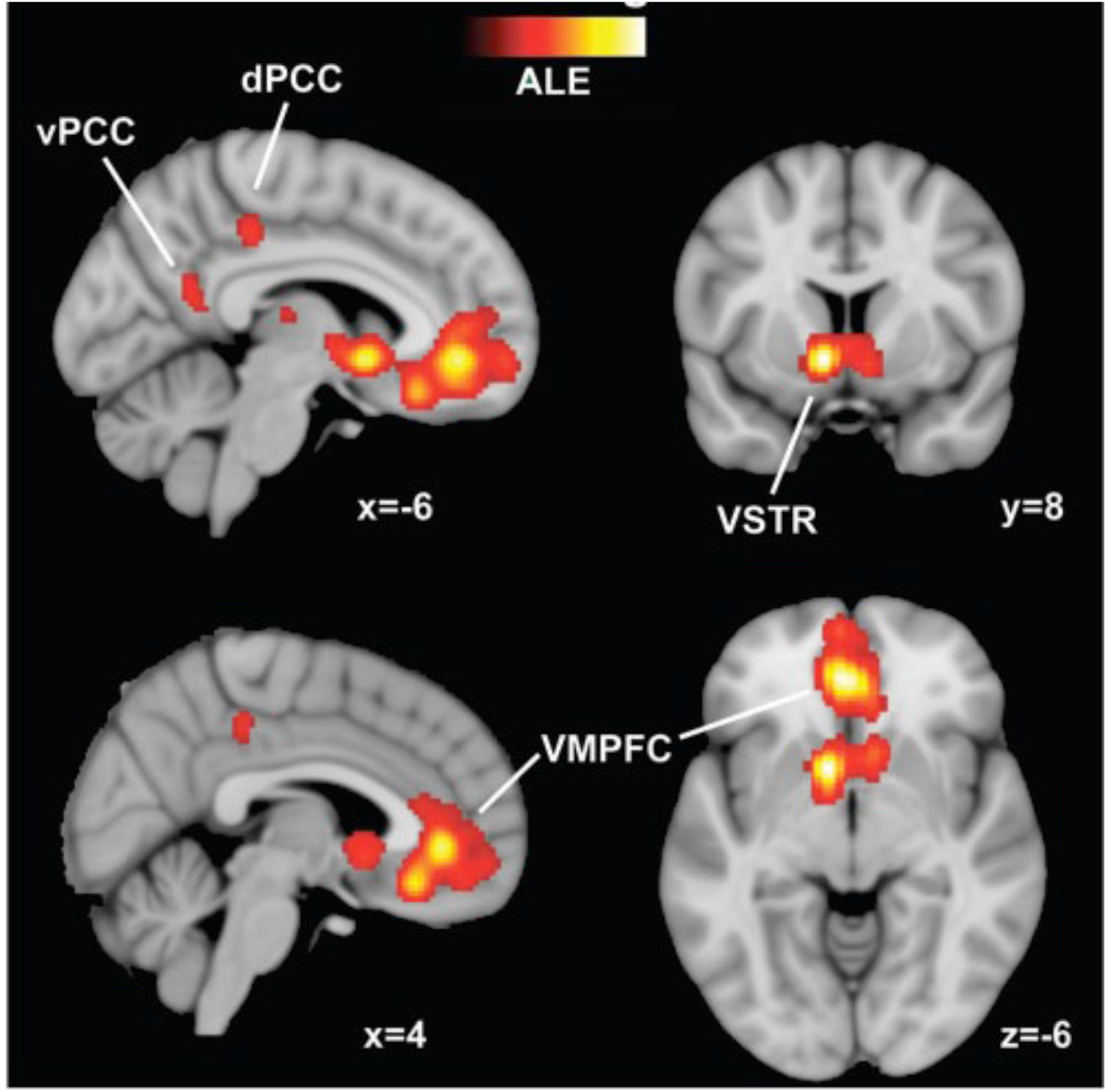
Representation of Value in the Human Brain. *Notes*. Regions of the brain that represent value, identified via a meta-analysis of neuroimaging studies. ALE = Activation Likelihood Estimation (brighter regions indicate greater signal strength across studies included in the meta-analysis); dPCC = dorsal posterior cingulate cortex; vPCC = ventral posterior cingulate cortex; VSTR = ventral striatum; VMPFC = ventromedial prefrontal cortex. Reproduced with permission from Clithero and Rangel (2014).

This body of evidence has led to the common-currency hypothesis, which suggests that a small set of specific brain areas appears to encode the subjective values associated with many different types of rewards on a common neural scale, regardless of the variation in the stimulus types giving rise to the evaluations (Levy & Glimcher, 2012). Perhaps not surprisingly, the same set of regions also underlies aesthetic experiences, given that our preferences for attractive faces (Kim et al., 2007; O’Doherty et al., 2003), harmonious color combinations (Ikeda, Matsuyoshi, Sawamoto, Fukuyama, & Osaka, 2015), geometrical shapes (Jacobsen, Schubotz, Höfel, & Cramon, 2006), and paintings or musical excerpts (Ishizu & Zeki, 2011) also reflect the extent to which we assign subjective value to stimuli of varying reward properties (see also Brown, Gao, Tisdelle, Eickhoff, & Liotti, 2011; Salimpoor & Zatorre, 2013; Vartanian & Skov, 2014). Moreover, functional connectivity between the nucleus accumbens and vmPFC predicts how much participants are willing to spend on musical excerpts (Salimpoor et al., 2013), suggesting that evaluative processes can also influence economic choices. These findings suggest that the brain networks supporting subjective valuation are also implicated in aesthetic judgements. We will argue below that this involvement extends to creative cognition.

Based on findings from neuroeconomics and studies of preference formation, we can advance a new conceptualization of creativity. Specifically, previous work suggests that two key processes support creative cognition: generation and evaluation of ideas (Basadur et al., 1982; Campbell, 1960; Simonton, 1999, 2013, in press). Generation involves coming up with many possible kernel solutions or ideas in response to a problem (or prompt), whereas evaluation refers to testing those solutions and ideas and selecting the best option(s) available. Here we posit that these processes also involve constructing and assessing the subjective value of ideas by integrating different criteria or attributes such as novelty and usefulness (Runco & Jaeger, 2012; Sternberg, 1999). Thus, the current framework proposes that value-based decision-making processes (e.g., value assignment, representation, comparison) underlie creative cognition.

Value-based decision-making models assume that choices are made by assigning an overall value to each option, computed as the weighted sum and/or product of different attributes (e.g., Hutcherson, Bushong, & Rangel, 2015; Hutcherson, Montaser-Kouhsari, Woodward, & Rangel, 2015; Suzuki, Cross, & O’Doherty, 2017). For example, neurocomputational evidence suggests that when making food decisions, the value of a food is dynamically constructed from the weighted sum of two attributes: perceived health and taste (Hare et al. 2009; Sullivan, Hutcherson, Harris, & Rangel, 2015). Whether an individual chooses to consume a healthy or unhealthy food (e.g., chips, broccoli) depends on not just the perceived health and taste of the food, but also the weights assigned to each attribute, which can be modulated by contextual factors (Hare, Malmaud, & Rangel, 2011). The specific decision context determines which attributes will be considered, as well as the strength of the weights assigned to each attribute (e.g., value of helping might depend on the weighted sum of how much one cares about oneself and others; see Hutcherson et al., 2015).

We suggest that within the context of creativity, the value of an idea will also be dynamically constructed from the weighted sum and/or product of attributes such as novelty and usefulness and/or their interaction. Because the weights assigned to each attribute change in different contexts, novelty and usefulness might not contribute to overall subjective value to the same extent across all contexts. Consistent with these ideas, Diedrich et al. (2015) found that judgments of usefulness only come into play after an idea has been deemed novel, suggesting that the weights assigned to each attribute might change at different stages of evaluation. While the field has focused primarily on the attributes of novelty and usefulness, our framework is not limited to these attributes. Indeed, we hope to provide a general framework for investigating how other attributes and/or contextual factors could also contribute to the computation of value in different contexts (e.g., surprise, see Simonton, 2012).

Conceptualizing creative cognition as value-based decision-making leads to several novel neurobiological predictions. First, it predicts that computations in neuroeconomic valuation regions of the brain (e.g., mPFC, OFC, PCC) should be associated with evaluative processes during creative cognition. Indeed, this prediction has already found support in fMRI studies that explicitly compared generative and evaluative processes during creative cognition. For example, Ellamil, Dobson, Beeman and Christoff (2012) focused on creative drawing, and instructed participants in the fMRI scanner to first design book covers and then to subsequently evaluate their designs and ideas. Compared to the generation of drawings, their evaluation was associated with greater activation in a set of regions including the medial frontal gyrus and PCC—both of which are involved in value-based decision making. Similarly, Mayseless, Aharon-Peretz, and Shamay-Tsoory (2014) demonstrated that evaluating the originality of ideas was associated with activation in a set of regions including the PCC. Further, a recent electroencephalogram (EEG) study found that evaluating ideas improved originality on a divergent thinking task, and was associated with increased frontal alpha activity that might reflect memory retrieval and integration processes (Hao et al., 2016). Neuroeconomic EEG studies have also shown that valuation processes reflect the integration of information from sensory and memory-related regions (Harris, Adolphs, Camerer, & Rangel, 2011). Together, these results support our first prediction, and underscore the role played by value-based decision-making processes during idea evaluation. They also indicate that EEG can be used to examine the temporal dynamics of valuation processes during creative cognition.

Second, because increased fMRI BOLD activity in valuation regions has been associated with increased subjective value (e.g., Kable & Glimcher, 2007), we also predict that neural responses in those regions should correlate positively with the perceived creativeness of ideas (weighted sum and/or product of novelty and usefulness) generated during creative cognition. For example, when performing divergent thinking tasks such as the alternate uses task, participants’ self-reported ratings of originality for their responses should correlate positively with activity in regions such as the mPFC, OFC, and PCC. Finally, given that neural responses in these valuation regions can predict economic choices (Smith, Bernheim, Camerer, & Rangel, 2014; Tusche, Bode, & Haynes, 2010), it might be possible to use these neural responses (combined with machine learning; see Shrivastava, Ahmed, Laha, & Sankaranarayan, 2017) to predict which idea, out of all the ideas that have been generated, will be selected eventually by the individual as the best idea.

### What makes something creative?

Neuroeconomics also seeks to develop neurologically plausible computational models that specify which decision variables (e.g., attributes) are computed, how they are computed in distinct brain regions and networks, and how these computations lead to choices (Rangel & Hare, 2010; Ratcliff, Smith, Brown, & McKoon, 2016; Shadlen & Kiani, 2013; Smith & Ratcliff, 2004). These models have proven fruitful in various domains such as perceptual decision making (Churchland, Kiani, & Shadlen, 2008; Gold & Shadlen, 2007), memory (Shadlen & Shohamy, 2016), self-control (Berkman, 2017; Berkman, Hutcherson, Livingston, Kahn, & Inzlicht, 2017; Hare, Camerer, & Rangel, 2009), and social decisions (Ruff & Fehr, 2014). We believe a neuroeconomic framework can also be useful for explaining creative cognition.

The assumptions underlying most neurocomputational models are that a noisy relative value signal accumulates over time, and that decisions are made once the accumulated information or evidence for one option becomes sufficiently strong to drive choice. For example, one study showed that individuals make altruistic or selfish choices by assigning an overall value to each option—computed as the weighted sum of two attributes: reward for self and reward for the other person (Hutcherson et al., 2015). Importantly, information about the two attributes were computed independently in distinct brain regions before being integrated and represented as an overall value signal in the vmPFC, and these processes could be described using extensions of standard computational models of decision making (e.g., Ratcliff et al., 2016). Given that creative judgements or valuations are assumed to depend on the integration of multiple attributes, in what follows we will outline how neurocomputational models may provide insights into attribute integration during creative cognition.

The assumptions underlying multi-attribute integration computational models of choice resemble those made in models of aesthetic experiences. Chatterjee and Vartanian (2014, 2016) suggested that distinct neural systems underlie different aspects of aesthetic experiences (e.g., emotional, perceptual), and that different weights might be assigned to different systems that underlie those aspects. For example, studies have shown that humans prefer curved over sharp objects (Bar & Neta, 2006), and that sharp objects tend to increase activity in the amygdala (Bar & Neta, 2007), presumably reflecting increased arousal, salience, or sense of threat associated with sharp objects. Neurocomputational models would thus predict that activity in the amygdala might reflect one of the many attributes (e.g., sense of threat) that an individual might consider when computing the overall liking for a sharp or curved object (computed within the brain’s valuation system). Because creative ideas are also defined along multiple attributes, future work could explore how information about different attributes (and perhaps beyond just novelty and usefulness; e.g., surprise) are computed and weighed in distinct brain regions, and how their interaction and integration causes an individual to evaluate an idea or product high or low on creativity based on its subjective value. This proposal is consistent with Martindale’s (1984) theory of cognitive hedonics, according to which thoughts (e.g., ideas) have evaluative aspects, which in turn can drive one’s preference for and continued pursuit of certain ideas over others. If the common currency hypothesis were correct (e.g., Levy & Glimcher, 2012), then the evaluation of ideas should also occur within the same neural network that computes subjective values for all other stimuli.

## Locus coeruleus-norepinephrine (LC-NE) system supports creative cognition

### Exploiting and exploring ideas

When trying to generate creative ideas, we assign higher subjective value to ideas that are high in terms of both novelty and usefulness. In cases where that occurs, it is often advantageous to exploit the idea further. In contrast, if an idea has relatively low subjective value (i.e., it is low in novelty and usefulness), it may be preferable to explore other ideas to find better alternatives. Many decisions in our daily lives require us to make a trade-off between exploitation and exploration (Christian & Griffiths, 2016; Cohen, McClure, & Yu, 2007; Hills, Todd, Lazer, Redish, & Couzin, 2015). Do we search for new ideas or stick with existing ones? Which ideas should you pursue? For example, when inventing a new product, should you continue developing new ideas after having generated *n* number ideas or should you start to focus on and develop one of them further? How do our brains choose the best course of action—or the best creative solution?

Our framework suggests that activity in the locus coeruleus-norepinephrine (LC-NE) neuromodulatory system plays an essential role in creative cognition by modulating the balance between exploitation and exploration. Our framework focuses mainly on the LC-NE system, but we hasten to note that all the major neuromodulatory systems have also been implicated in various decision and valuation processes that underlie creative cognition (e.g., Spee, Ishizu, Leder, Mikuni, Kawabata, & Pelowski, in press). For example, the dopamine system may be necessary for learning which ideas are rewarding or creative because dopamine is believed to be important for learning the value of objects from prediction errors (Berke, 2018; Montague, Hyman, & Cohen, 2004; Roesch, Calu, & Schoenbaum, 2007; Schultz, 2007). Serotonin, like dopamine, has also been implicated in reward signaling—specifically learning from punishments or negative prediction errors (Boureau & Dayan, 2011; Cools, Robinson, & Sahakian, 2008; Cools, Nakamura, & Daw, 2011; Kranz, Kasper, & Lanzenberger, 2010; Nakamura, Matsumoto, & Hikosaka, 2008). Moreover, acetylcholine and norepinephrine appear to play major roles in flexible learning and decision making; especially relevant to our framework is that norepinephrine has been proposed to mediate flexible shifts between exploitation and exploration (Aston-Jones & Cohen, 2005b, Kehagia, Murray, & Robbins, 2010; Yu & Dayan, 2005). If creative cognition were mediated by processes that resemble those in classic exploitation-exploration trade-offs (Cohen et al., 2007; Daw, O’Doherty, Dayan, Seymour, & Dolan, 2006), then understanding the relationship between decision making and creative processes will organize and benefit research in various fields (we address the interplay between various neuromodulatory system further under *Interactions with other neurotransmitter systems*).

Creative cognition appears to rely on our abilities to exploit and explore ideas, as well as switch between these two modes of cognition (Monechi, Ruiz-Serrano, Tria, & Loreto, 2017). When people are initially trying to find inspiration or ideas for tackling a new problem, they are attempting to explore and generate ideas that satisfy certain criteria that are often based on relatively abstract goals. The relative importance of each criterion depends on the context, and the subjective value of an idea will depend on how well it satisfies those criteria. For example, an artist might be seeking an idea that best conveys a particular meaning, and a scientist might be developing a new experimental procedure that most stringently tests a theoretical prediction. These individuals are generating ideas by exploring the available options and pruning them by assessing their subjective values. Different ideas will have different subjective values depending on how well they satisfy certain criteria. Most ideas will likely be entertained very briefly because they fail to satisfy those criteria (i.e., they will be assigned low subjective values), and are subsequently discarded. However, when individuals land on an idea that satisfies those criteria sufficiently, they will likely stop exploring alternatives because they would want to devote their time and resources to fully exploit the value it provides. The present framework suggests that the creative process described above reflects an adaptive value-optimization process mediated by activity in the LC-NE system and interconnected brain regions that compute and evaluate the subjective values of various creative ideas and strategies (see Aston-Jones & Cohen, 2005b), while at the same time also acknowledging that other neuromodulatory systems likely also contribute to the dynamics of creative cognition.

### Locus coeruleus-norepinephrine system and function

The locus coeruleus nucleus sits deep in the pons and sends noradrenergic projections to nearly all brain regions (with the notable exception of the basal ganglia and hypothalamus). It is also the only source of norepinephrine (also known as noradrenaline) to the cerebral, cerebellar, and hippocampal cortices (Foote & Morrison, 1987; Moore & Bloom, 1979) (Figure 2). Because the locus coeruleus projects diffusely to cortical regions, early research focused primarily on its role in general cognitive processes, especially in mediating arousal (Amaral & Sinnamon, 1977; Berridge & Waterhouse, 2003).

**Figure 2.**
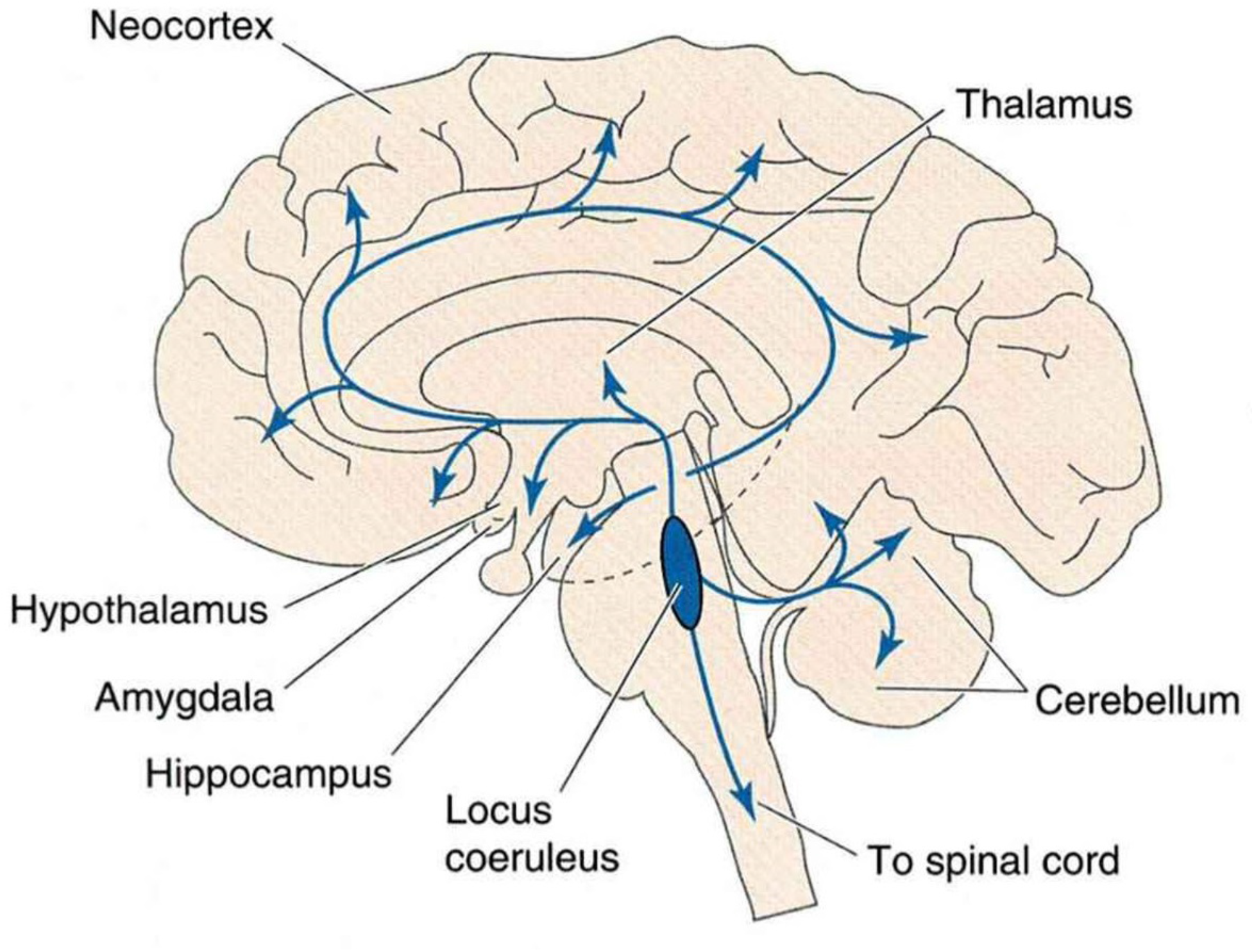
The Locus Coeruleus-Norepinephrine (LC-NE) System. *Notes*. Reproduced with permission from Breedlove, Watson, and Rosenzweig (2010).

Recent work has highlighted the role of the LC-NE system in regulating engagement during tasks that require selective or focused attention (Aston-Jones & Cohen, 2005a-b; Chmielewski, Mückschel, Ziemssen, & Best, 2017). Many studies have shown that salient or task-relevant stimuli reliably elicit *phasic* activation of locus coeruleus neurons and norepinephrine release in cortical target sites (Aston-Jones & Bloom, 1981; Hervé-Minvielle & Sara, 1995). Phasic activity is characterized by short-duration, rapid bursts of locus coeruleus activity and concomitant norepinephrine release whose timing correlates strongly with behavioral performance (Usher, Cohen, Servan-Schreiber, Rajkowski, & Aston-Jones, 1999). Such phasic activity is thought to increase neural gain (sensitivity or responsivity) in task-relevant cortical regions, which then filter task-relevant events to improve engagement and performance (Aston-Jones & Cohen, 2005b; Hasselmo, Linster, Patil, Ma, & Cekic, 1997; Lee et al., 2018; Mather, Clewett, Sakai, & Harley, 2016).

While phasic activity is often tightly coupled with good performance during selective attention tasks, *tonic* activity also affects engagement and performance (Aston-Jones & Cohen, 2005b). In contrast to phasic activity, tonic activity is characterized by intrinsic, ongoing locus coeruleus firing that is unrelated to the task and occurs in the background. Specifically, relative levels of phasic and tonic locus coeruleus activity relate to performance in a manner that reflects the classic Yerkes-Dodson inverted-U arousal curve (Yerkes & Dobson, 1908): At moderate levels of tonic activity, phasic activity is elevated and performance is optimal, whereas shifts towards relatively lower or higher tonic activities are associated with reduced phasic activity and poorer performance on selective attention tasks. More broadly, the *phasic mode* is characterized by moderate tonic activity that facilitates exploitation of options, whereas the *tonic mode* is characterized by relatively higher tonic activity that is thought to promote exploration of alternatives that could be potentially more rewarding (Aston-Jones & Cohen, 2005b; Berridge & Waterhouse, 2003; Cohen et al., 2007; Usher et al., 1999).

Here we build on evidence from selective attention tasks and neuroeconomics to provide an integrative neurobiological framework for creative cognition. The present framework suggests that creative cognition relies on neural value computations that also underlie a range of behaviors such as choices between material goods, perceptual decisions, memory, and social decisions (e.g., Berkman et al., 2017; Ruff & Fehr, 2014; Shadlen & Kiani, 2013). Drawing on recent theorizing on role of the norepinephrine system (Aston-Jones & Cohen, 2005b; Kehagia et al., 2010, Sadacca, Wikenheiser, & Schoenbaum, 2017), we suggest that value-based decision-making processes mediate the fine balance and transitions between the phasic and tonic locus coeruleus modes, which, in turn optimize creativity by facilitating flexible exploitation and exploration of ideas and strategies with varying subjective values. That is, flexible and adaptive fluctuations between phasic and tonic modes may be critical to creative cognition, and might underlie performance on both laboratory and real-world measures of creativity.

### Norepinephrine underlies creative cognition

Being creative depends on one’s ability to both maintain task goals (e.g., exploit specific mental representations of ideas or familiar strategies) as well as switch between task sets (e.g., explore different mental representations (see Goschke, 2000; Hills et al., 2015; Monsell, 2003). Thus, like many everyday value-based decision dilemmas, the demands on creative cognition seem to imply a delicate balance between exploitative and exploratory processes that are regulated by the phasic and tonic locus coeruleus modes, respectively. Indeed, early work had already hinted at potential relationships between LC-NE activity, engagement, arousal, and creativity. For example, classic studies have shown that relaxed and less aroused states are associated with increased creativity (Martindale & Greenough, 1973). During creative generation, more creative individuals show stronger EEG alpha band activity (Martindale & Hasenfus, 1978), which is believed to reflect reduced arousal mediated by norepinephrine from the locus coeruleus (Foote, Berridge, Adams, & Pineda, 1991; Foote & Morrison, 1987).

Pharmacological studies have provided stronger evidence for the role of norepinephrine in creative processes (Beversdorf, 2013, in press; Heilman, 2016; Heilman, Nadeau, & Beversdorf, 2003). One such process is cognitive flexibility, required in set-shifting tasks where attention must be shifted from one perceptual dimension to another (Birrell & Brown, 2000). Increasing tonic norepinephrine via α-adrenergic receptors in the mPFC improved rats’ set-shifting performance (Lapiz, Bondi, & Morilak, 2007; Lapiz & Morilak, 2006), presumably because elevated tonic norepinephrine corresponds to a “scanning attention” mode that reduces focus on well-established cues or strategies and promotes attention to novel or previously non-salient cues (Aston-Jones, Rajkowski, & Cohen, 1999; Sadacca et al., 2017). However, the effects of pharmacological interventions might depend on other factors, including personality. For example, administering methylphenidate (a psychostimulant that increases norepinephrine and dopamine levels in the brain) improves creativity only in people low in novelty seeking (i.e., weaker exploratory tendencies), but impairs creativity in those high in novelty seeking (Gvirts et al., 2017). These findings suggest that when designing and interpreting norepinephrine (intervention) studies, we should consider contextual and personality variables that could influence the balance between phasic and tonic locus coeruleus modes (e.g., stress, age; see also Mather & Harley, 2016).

Although increasing norepinephrine via α-adrenergic receptors facilitates set shifting, reducing norepinephrine via *β*-adrenergic receptors (e.g., *β*-adrenergic blocker propranolol) seems to benefit other forms of cognitive flexibility that require efficient access to and search throughout neural networks (e.g., anagram problems; Beversdorf, Hughes, Steinberg, Lewis, & Heilman, 1999; Beversdorf, White, Chever, Hughes, & Bornstein, 2002; Campbell, Tivarus, Hillier, & Beversdorf, 2008; Hecht, Will, Schachtman, Welby, & Beversdorf, 2014). These opposing effects suggest that *α*- and *β*-adrenergic receptors might mediate creative cognition via distinct processes, possibly owing to differences in whether the specific receptor in question is excitatory or inhibitory, as well as their differential affinities for norepinephrine. For example, compared with *α*-2A receptors, *β*-receptors tend to have lower affinity and activate only with relatively high levels of norepinephrine (Lee et al., 2018; Samuels & Szabadi, 2008). Thus, to better understand how phasic and tonic norepinephrine modes mediate creative cognition, future research should investigate and compare the effects of both activating and blocking *α*- and *β*-adrenergic receptors in contexts that have been associated with different levels of tonic norepinephrine (e.g., drowsiness, wakeful alertness, and stress are associated with low, moderate, and high norepinephrine levels respectively).

For example, stress is known to increase norepinephrine, impair prefrontal function, and alter cellular activity in prefrontal norepinephrine neurons (Arnsten, 2009; Miner, Jedema, Moore, Blakely, Grace, & Sesack, 2006; Morilak et al., 2005; Goldfarb, Froböse, Cools, & Phelps, 2017). Critically, the benefits of administering *β*-adrenergic blockers on certain forms of cognitive flexibility are most apparent under stressful conditions (Alexander, Hillier, Smith, Tivarus, & Beversdorf, 2007). These findings again suggest that whether pharmacological interventions improve or impair creativity might depend on factors (e.g., personality, context) that affect tonic norepinephrine levels.

While the studies described thus far have demonstrated the broad causal role of norepinephrine on exploitation, exploration, and creative cognition, they have not shown how specifically phasic and tonic locus coeruleus activities during creative cognition are affected by manipulating norepinephrine levels. Moreover, these studies often focused on cognitive flexibility in the context of situational stress (e.g., Alexander et al., 2007; Campbell et al., 2008). This contextual factor is particularly relevant because stress could have increased tonic norepinephrine levels, shifted the baseline balance between exploitation and exploration tendencies, influenced attention and performance, and engaged *α*- and *β*-receptors differently (Arnsten, 2000, 2009; Berridge & Waterhouse, 2003; Robbins & Arnsten, 2009). Moreover, as natural phasic activity may be influenced by general changes in tonic activity (e.g., stress, receptor agonists), this phasic-tonic duality in firing modes often complicates the interpretation of many pharmacological studies (discussed in the context of dopamine by Beninger & Miller, 1998). Nevertheless, these studies have provided strong evidence that the balanced fluctuations in norepinephrine levels are essential to different forms of cognitive flexibility. Future work should seek to resolve the disparate findings of why sometimes increasing norepinephrine benefits creative processes, and other times reducing norepinephrine benefits these processes. Examining the neurobiological processes and contextual factors that determine changes in norepinephrine levels and transitions between the two locus coeruleus modes will also be crucial to understanding how norepinephrine underlies exploit-explore trade-offs during creative cognition (e.g., Aston-Jones & Cohen, 2005b).

### Value-based decision making regulates transitions between exploitation and exploration

The adaptive gain theory of LC-NE function (Aston-Jones & Cohen, 2005b) may help bridge neuroeconomic findings with research relating norepinephrine to creativity. According to this theory, during selective attention tasks, the high subjective value associated with the current task triggers the phasic mode, which facilitates exploitation of ongoing behaviors or existing strategies to optimize performance. Low or declining subjective value, however, triggers the tonic mode, which promotes disengagement from the current task and exploration of alternatives that can potentially represent more rewarding opportunities (Aston-Jones & Cohen, 2005b; Cohen et al., 2007).

The present framework suggests that the phasic locus coeruleus mode corresponds to creative processes that involve exploitation of ideas or strategies with high subjective values. For example, when solving anagrams (a standard measure of cognitive flexibility) in an experiment, an idea or strategy that allows one to generate many solutions within a limited time is likely to have relatively high subjective value. One common anagram strategy involves finding suffixes and adding them to the end of existing words (e.g., paint-er, work-*er*). Value-based decision processes will then trigger the phasic mode that helps exploit this high-value solution through processes like evaluation or elaboration (e.g., find additional suffixes: paint-ing, work-ing). Thus, at least for laboratory measures of creativity, the phasic mode should increase neural gain and filter task-relevant representations to help generate solutions, like how it facilitates good performance during selective attention tasks (Aston-Jones & Cohen, 2005b; Mather et al., 2016).

The tonic mode, however, is associated with processes that facilitate the exploration of alternatives when the subjective values of existing options are relatively low or declining. When an idea or strategy is no longer novel or successful in generating novel and useful solutions, the declining subjective value triggers relative shifts towards the tonic mode that promotes exploration of alternatives. Although this mode might temporarily impair immediate performance by causing increased distractibility and temporary disengagement from the currently salient task representations (e.g., finding suffixes), it encourages individuals to widen their attentional focus to explore alternatives that might provide better long-term payoff despite the short-term costs of exploration (Aston-Jones & Cohen, 2005b; Sadacca et al., 2017; Tervo et al., 2014; Usher et al., 1999). For example, if one has exhausted all possible suffixes for a given anagram, the shift from phasic to tonic mode might trigger changes in strategies, causing one to start considering prefixes instead and adding them to the beginning of existing words (e.g., *re*-paint, *re*-work). Thus, the tonic mode might be required—at least temporarily—to mentally explore alternatives. Moreover, because creative cognition appears to resemble basic ecological problems the brain has evolved to solve in real-world environments (e.g., exploitation-exploration dilemmas, animal foraging in patchy environments; Cohen et al., 2007; Kidd & Hayden, 2015; Mobbs, Trimmer, Blumstein, & Dayan, 2018; Pearson et al., 2014), this framework also has potential to explain not only creativity in the laboratory, but also real-world creativity.

Critically, the adaptive gain theory suggests that whether LC-NE activity is in the phasic or tonic mode depends on value computations in cortical regions such as the OFC (Padoa-Schioppa & Assad, 2006) and the anterior cingulate cortex (ACC; Calhoun & Hayden, 2015; Heilbronner & Hayden, 2016; Shenhav et al., 2013)—both of which project densely to the locus coeruleus (Aston-Jones & Cohen, 2005a; Porrino & Goldman-Rakic, 1982). The present framework suggests that during creative cognition, activity in the neural valuation regions (e.g., OFC) drives and produces the transitions between the phasic and tonic locus coeruleus modes.

When a newly generated idea or strategy is novel and useful, the valuation regions would assign a high subjective value to it, triggering the phasic mode that promotes exploitation of that idea. But when ideas are not or no longer useful or novel, the valuation regions would register low overall subjective value, which temporarily triggers relative shifts towards the tonic mode that increases baseline norepinephrine release, facilitating exploring and sampling of other ideas that might provide higher long-term subjective value (Aston-Jones & Cohen, 2005b). In sum, the subjective values assigned to ideas or strategies (determined by how well they satisfy criteria such as novelty and usefulness) are hypothesized to flexibly balance the transitions between the phasic and tonic modes. These transitions, in turn, help maximize long-term payoff by optimizing the trade-off between exploitation and exploration.

### Reinterpretation of existing findings, new predictions, and new measures

By extending the adaptive gain theory of LC-NE function to creative cognition, the present framework not only helps to reinterpret and integrate existing findings, but also makes new predictions that can be tested with various behavioral and neurophysiological measures. First, because the LC-NE system is hypothesized to drive exploitation-exploration processes that underlie creative cognition, one would expect the phasic and tonic locus coeruleus modes to respectively correspond to exploiting ideas with high subjective value (e.g., evaluation and/or elaboration) and exploring alternative options (e.g., switching to a different strategy). These predictions can be tested with laboratory measures of creativity by tracking participants’ behavioral performance and strategy use while measuring fMRI BOLD activity in the locus coeruleus and valuation regions (Kolling, Behrens, Mars, & Rushworth, 2012; Murphy, O’Connell, O’Sullivan, Robertson, & Balsters, 2014).

In addition, measures such as pupil diameter can be used to study how creative processes unfold in real time because they track LC-NE activity and elucidate the processes underlying value-based decision making (Hassall, Holland, & Krigolson, 2013; Lin, Saunders, Hutcherson, & Inzlicht, 2018; Murphy, Robertson, Balsters, & O’Connell, 2011; Van Slooten, Jahfari, Knapen, & Theeuwes, 2018). For example, phasic LC-NE activity correlates with the P3, a positive potential that peaks about 350 ms following stimulus onset and observed over central-parietal midline EEG electrodes (Nieuwenhuis, Aston-Jones, & Cohen, 2005); changes in locus coeruleus firing rates also correspond remarkably well to changes in pupil dilation responses (Joshi, Kalwani, & Gold, 2016; Murphy et al., 2014; Reimer et al., 2016; Varazzani et al., 2015). Moreover, in gambling tasks designed to specifically investigate exploitation-exploration trade-offs, high baseline pupil diameter (elevated tonic activity) has been shown to predict disengagement and exploration of alternative rewards, whereas low baseline pupil diameter predicted task engagement and exploitation of the current reward (Jepma & Nieuwenhuis, 2011). Since the present framework proposes that value computation and exploitative-exploratory processes also underlie creative cognition, incorporating EEG and pupillometry in future research may provide insights into creative cognition and related processes (see Smallwood, Brown, Baird, Mrazek, Franklin, & Schooler, 2012; Unsworth & Robison, 2016; van der Wel & van Steenbergen, 2018). For example, one could pharmacologically manipulate norepinephrine and then track changes in tonic and phasic pupil diameter while participants perform cognitive flexibility tasks. Together, these findings and suggestions further highlight the usefulness of the present framework by proposing different measures that can be used to investigate the processes participants rely on while engaging in creative cognition (e.g., pupil diameter, locus coeruleus BOLD activity, P3; see also heart rate variability, Mather et al., 2017).

Second, the present framework suggests that instead of focusing solely on behavioral outcome measures such as response latency and/or responses generated during creativity tasks (e.g., fluency, defined as the number of ideas), we can gain more insights into creative cognition if we were to also investigate the underlying processes that lead to the observed outcomes. For example, although two individuals may have generated the same number of responses on a cognitive flexibility task, to what extent can we infer that they have relied on the same strategies and underlying processes to arrive at those solutions? For example, they might have relied differentially on exploitative and explorative strategies, despite having generated the same number of responses. By focusing on the underlying processes, our framework could explain why laboratory measures of creativity (e.g., divergent thinking) at times correlate weakly with real-world creative achievement (e.g., Zabelina, Saporta, & Beeman, 2016), which is often measured using the Creative Achievement Questionnaires (Carson, Peterson, & Higgins, 2005). Laboratory measures of creativity often require participants to generate many solutions within a limited amount of time. This emphasis on responding under time pressure in fact characterizes the demands of selective attention tasks, which are best met by the phasic locus coeruleus mode (Aston-Jones & Cohen, 2005b; Usher et al., 1999). Real-world creative achievement, however, is often more protracted (i.e., involves less immediate time pressure), and might ultimately require different dynamics than laboratory-based creativity tasks. In addition, the criteria defining the “correctness” of any given solution or idea might be relatively unclear and could even change over time. In this sense, discovering and stumbling upon better alternatives through exploration (triggered by the tonic mode) might be a particularly apt characterization of real-world creativity (Monechi et al., 2017). Consistent with these ideas, divergent thinking in the laboratory has been associated with selective attention, whereas creative real-world achievement has been associated with wider attentional focus and failures to inhibit seemingly irrelevant stimuli (Carson, Peterson, & Higgins, 2003; Zabelina, Colzato, Beeman, & Hommel, 2016; Zabelina, O’Leary, Pornpattananangkul, Nusslock, & Beeman, 2015; Zabelina et al., 2016). More broadly, creativity in laboratory and real-world tasks might be predicted by distinct patterns of exploitation-exploration tendencies, and these ideas will have to be tested in future experiments.

Third, people with greater real-world creative achievement appear to have wider attentional focus, which can in turn distract them from their primary tasks. Although distractibility usually impairs task performance, it might allow individuals to consider and generate more alternative ideas (e.g., Carson et al., 2003; Zabelina et al., 2015, 2016), and might be associated with increased exploratory tendencies that are driven by relatively higher tonic locus coeruleus mode activity and norepinephrine levels (but see below for stress and psychological dysfunction). Our framework therefore has the potential to explain not only the neurobiological bases of creativity, but also individual differences in creativity. Although there is no direct evidence for the hypothesized relationship between tonic activity and creativity, the locus coeruleus has been associated with individual differences in cognitive function and abilities (Mather & Harley, 2016). Indeed, a recent study found that baseline pupil diameter (a proxy for tonic activity) correlates with intelligence (Tsukahara, Harrison, & Engle, 2016), which is a factor that predicts individual differences in creativity (Benedek, Jauk, Sommer, Arendasy, & Neubauer, 2014; Jauk, Benedek, Dunst, & Neubauer, 2013; Jauk, Benedek, & Neubauer, 2014; Nusbaum & Silvia, 2011). But given that blocking norepinephrine has also been shown to benefit certain types of creative processes (e.g., Alexander et al., 2007; Hecht et al., 2014), increased tonic activity and norepinephrine might benefit only specific forms of creativity in certain contexts.

Fourth, increased tonic locus coeruleus activity could predispose creative people to increased distractibility, primarily because higher tonic activity increases intrinsic background neural activity, reduces attentional selectivity, which in turn allows a wider range of seemingly irrelevant mental representations to be sampled (Eldar, Cohen, & Niv, 2013; Hasselmo et al., 1997; Usher et al., 1999). However, whether these effects lead to better or worse creativity may depend on which norepinephrine receptors are activated and the specific creative process under consideration (e.g., Alexander et al., 2007; Lapiz & Morilak, 2006). Nevertheless, these effects suggest that creative people may be more likely to experience sensory overstimulation because of their over-inclusive attention. Consistent with these predictions, many studies have reported that creative people tend to exhibit greater sensitivity to sensory stimuli. For example, compared to less creative people, they rate electrical shocks as being more painful and register higher amplitude skin potential responses to tones (e.g., Martindale, 1977, Martindale, Anderson, Moore, & West, 1996; Martindale & Armstrong, 1974). Presently, the precise relationship between phasic-tonic locus coeruleus activity, norepinephrine receptor types, and individual differences in creativity remains unclear, and we believe our framework could offer insights into the interplay among these variables, as well as why and how locus coeruleus activity and sensory overstimulation are related in laboratory and real-world studies.

Real-world creative achievement has also been associated with “leaky” attention, which was reflected in reduced sensory gating as indexed by the P50 event-related potential (Zabelina et al., 2015). These findings suggest that real-world creative achievers might be less able to filter irrelevant information—a filtering process mediated by the phasic locus coeruleus mode—and that such leaky sensory gating (mediated by the tonic locus mode) might be one of the processes that benefit creativity by focusing attention on more stimuli regardless of their immediate relevance (Mendelsohn & Griswold, 1964; Russell, 1976). In addition, creative people are also hypothesized to connect distantly related concepts or ideas more easily, presumably because the tonic mode increases noise and leaky sensory gating, allowing them to sample a wider range of stimuli and cues (Ansburg & Hill, 2003).

Further support for the relationship between the tonic mode and leaky sensory gating comes from recent work showing that pupil diameter reflects locus coeruleus-driven neural gain and sensory processing, such that higher gain (i.e., phasic mode) is associated with narrow attentional focus, whereas lower gain (i.e., tonic mode) is associated with broader attentional focus (Eldar et al., 2013; Eldar, Niv, & Cohen, 2016). Despite the evidence linking the tonic mode with creativity, it could be that creative people are also better at balancing and switching between the phasic and tonic modes. Our framework suggests that by incorporating valuation processes, we can better understand how creative people excel at switching between different modes of cognition in the service of creative problem solving, which remains one of the major open questions in the field (see Dorfman, Martindale, Gassimova, & Vartanian, 2008; Vartanian, 2009; Vartanian, Martindale, & Kwiatkowski, 2007).

Finally, since the present framework ascribes a central role to the LC-NE system, it follows that disturbances in the LC-NE system might affect creative cognition. For example, the LC-NE system has been implicated in highly overlapping sets of clinical disorders associated with either enhanced or impaired creativity (e.g., schizophrenia, bipolar disorder; Baas, Nijstad, Boot, & De Dreu, 2016; Kyaga et al., 2011; MacCabe, Sariasian, Almqvist, Lichtenstein, Larsson, & Kyaga, 2018; Simonton, 2014). Some evidence suggests that schizophrenic patients have increased locus coeruleus cell volumes (Marner, Søborg, & Pakkenberg, 2005), and type 1 (positive symptoms) schizophrenia has been associated with elevated norepinephrine and metabolites in the brain (Yamamoto & Hornykiewicz, 2004). These patients often show sensory gating deficits, in that they fail to filter out potentially irrelevant stimuli (Braff, Geyer, & Swerdlow, 2001; Braff, Greenwood, Swerdlow, Light, & Schork, 2008). Moreover, increasing tonic activity leads to sensory gating deficits in rats, whereas reducing tonic activity via *α*-adrenergic receptors prevented these deficits (Alsene & Bakshi, 2011). Together, these findings suggest that the LC-NE system underlies gating deficits and mental disorders, but whether and how it explains changes in creativity related to these abnormalities remain open questions. Similarly, hypersensitivity to environmental stimuli—as reflected in increased rates of food allergies, asthma, and autoimmune diseases—have also been observed in people with high intelligence (Karpinski, Kolb, Tetreault, & Borowski, 2017), a trait that has been associated with increased creativity (e.g., Benedek et al., 2014). Given these potential links between LC-NE activity, sensory gating, real-world creative achievement (Zabelina et al., 2015, 2016), and flexible decision making (Aston-Jones & Cohen, 2005b; Sadacca et al. 2017), we suggest that future research consider how value-based decision-making and LC-NE processes might explain the relationship between creativity and certain clinical disorders.

Viewing our framework from the perspective of Carson’s (2011, 2014, in press) shared-vulnerabilities model could also help to better understand the relationship between the LC-NE system, creativity, and psychopathology. According to this model, both creative people and those suffering from psychopathology share certain vulnerabilities, including novelty seeking, cognitive disinhibition, and neural hyperconnectivity. For example, both creative people and those with schizophrenia or schizotypy have been shown to exhibit lower levels of latent inhibition—defined as the ability to screen from current attentional focus stimuli previously experienced as irrelevant (see Carson et al., 2003, p. 499; see also Eysenck, 1995). If unchecked, reduced latent inhibition could lead to disturbances in cognition that are caused by a reduced ability to discriminate between task-relevant and task-irrelevant information. However, what distinguishes creative people from those suffering from psychopathology is the additional presence of protective factors among creative people, including high intelligence, working memory capacity, and ego strength. In turn, the presence of these protective factors enables creative people to utilize their vulnerabilities in the service of goal-directed behavior. For example, greater working memory capacity might enable a person to systematically use and combine stimuli previously experienced as irrelevant to generate creative solutions (e.g., De Dreu, Nijstad, Baas, Wolsink, & Roskes, 2012). Within this framework, individual differences in LC-NE system activity might interact with vulnerability and protective factors to modulate creativity. However, to the best of our knowledge, there is no direct evidence linking different psychopathologies to different levels of tonic and phasic LC-NE activity, and remains an open question for investigation.

### Interactions with other neurotransmitter systems

While we have focused on the LC-NE system, much evidence suggests that other neuromodulatory systems—especially dopamine—also support creative cognition (e.g., Spee et al., in press). For example, converging evidence suggests that moderate (but not low or high) levels of dopamine in the striatum and prefrontal cortex facilitate different types of creative processes (Boot, Baas, van Gaal, Cools, & De Dreu, 2017). Different dopamine receptor subtypes in the mPFC and genes (e.g., dopamine D2 receptor, COMT) have been associated with cognitive flexibility and divergent thinking (Floresco, Magyar, Ghods-Sharifi, Vexelman, & Tse, 2006; Reuter, Roth, Holve, & Hennig, 2006; Zabelina et al., 2016; Zhang, Zhang, & Zhang, 2014). In addition, given dopamine’s role in reward and reinforcement learning (O’Doherty, Cockburn, & Pauli, 2017; Schultz et al., 1997), it might also be critical to learning what is creative and which actions or strategies lead to greatest creativity. Serotonin genes have also been associated with creativity (Reuter et al., 2006; Volf, Kulikov, Bortsov, & Popova, 2009), and serotonin has been implicated in specific forms of cognitive flexibility (e.g., reversal learning) that are primarily mediated by the valuation region OFC (Clark, Dalley, Crofts, Robbins, & Roberts, 2004; Clarke, Walker, Crofts, Dalley, Robbins, & Roberts, 2005). Similarly, acetylcholine has been associated with reversal learning (Robbins & Roberts, 2007; see also Yu & Dayan, 2005). A discussion of the theories and functions of these systems is beyond the scope of this article, but the above findings, together with work indicating that dopamine and serotonin play major roles in learning and valuation (e.g., Boureau & Dayan, 2011; Cools et al., 2011; Montague et al., 2004; Schultz, 2007), is consistent with our suggestion that valuation processes may underlie creative cognition.

Norepinephrine assumes a central role in our framework because of its proposed role in mediating the balance between exploitation and exploration during creative cognition, but some evidence suggests that dopamine and serotonin also modulate the phasic and tonic locus coeruleus modes (e.g., McClure, Gilzenrat, Cohen, 2006). Specifically, tonic dopamine and serotonin levels have been proposed to track average levels of reward and punishment (Boureau & Dayan, 2011; Cools et al., 2011; Niv, Daw, Joel, & Dayan, 2007), which might in turn determine the threshold for exploring alternatives (Hills et al., 2015). For instance, higher average reward rates, reflected in relative increases in tonic dopamine, might increase phasic mode activity in the LC-NE system, which corresponds to exploitative behaviors such as faster and more vigorous responding (e.g., Hamid et al., 2016; Salamone & Correa, 2002). Further evidence for the role of dopamine in governing these behaviors comes from a genetic study, which found that the dopamine D2 receptor and COMT genes were associated with exploitation and exploration (Frank, Doll, Oas-Terpstra, & Moreno, 2009). Together, these findings suggest that conceptualizing creative processes as involving valuation and exploitation-exploration trade-offs can potentially elucidate the roles of not just norepinephrine, but also the other neurotransmitters during creative cognition.

## Valuation processes and LC-NE activity mediate creative cognition network dynamics

Recent neuroimaging work has converged on the view that creative cognition involves dynamic interactions within and between large-scale brain networks, especially the default mode network (DMN) and the executive control network (Beaty, Benedek, Kaufman, & Silvia, 2015; Ellamil et al., 2012; Liu et al., 2015). The DMN and executive control network are engaged by different types of tasks. Specifically, the DMN is activated by tasks that involve internally-directed processes such as self-generated thought, future event simulation, and spontaneous thought, and it exhibits decreased activation during tasks that involve attention to external stimuli (Andrews-Hanna, Smallwood, & Spreng, 2014; Christoff, Irving, Fox, Spreng, & Andrews-Hanna, 2016; Mittner, Hawkins, Boekel, & Forstmann, 2016; Smallwood & Schooler, 2015; Zabelina & Andrews-Hanna, 2016). In contrast, the executive control network is part of a “task positive” set of regions that exhibit greater activation during tasks that require attention to external stimuli (Dixon et al., 2017). The observation of their joint activation during creative cognition has led to the idea that the two networks support different aspects of creativity: whereas the DMN supports creative idea generation, the executive control network modulates activity in the DMN to ensure that task goals are met (Beaty, Benedek, Silvia, & Schacter, 2016; Beaty et al., 2015). Although these brain networks are clearly implicated in creative cognition, one critical question remains unaddressed: What determines the engagement of these networks, as well as the interactions and transitions between them? The present framework speculates that network dynamics observed during creative cognition are driven by value computations in regions within the brain’s valuation system (Figure 1) and activity in the LC-NE system (Figure 2), which jointly optimize the trade-off between idea exploitation and exploration.

The core brain regions that assign, represent, and evaluate subjective value during value-based decision making are the OFC, vmPFC, and PCC (Bartra et al., 2013; Clithero & Rangel, 2014). Coincidentally, the mPFC and PCC also form the core of the DMN (Andrews-Hanna, Reidler, Sepulcre, Poulin, & Buckner, 2010; Zabelina & Andrews-Hanna, 2016). These anatomical (and related functional) overlaps suggest that DMN activity might in part reflect neural value computations proposed to underlie creative cognition in the present framework. For example, multiple lines of work suggest that the PCC might play an important role during creative cognition. Apart from being implicated in value-based decision making (Barack, Chang, & Platt, 2017; Bartra et al., 2013; Grueschow et al., 2015) and internally-oriented cognition (Zabelina & Andrews-Hanna, 2016; Christoff et al., 2016), the PCC might mediate functional coupling and transitions between different brain networks. For example, during early phases of divergent thinking, the PCC strongly couples with the salience network regions such as the insula, whereas during later phases it couples with executive control network regions (e.g., dorsal lateral PFC; Beaty et al., 2015). A primary function of the salience network is to focus the spotlight of attention on relevant stimuli in the service of goal-directed behavior, as well as initiating the switch between the DMN and the executive control network (Cocchi, Zalesky, Fornito, & Mattingley, 2013; Menon, 2015; Uddin, 2015). These findings suggest that in conjunction with nodes within the salience network, computations in the PCC might be critical for engaging different brain networks, as well as mediating network interactions and transitions in the service of creativity.

Indeed, neuroeconomic studies have demonstrated that the PCC mediates shifts between networks and corresponding transitions in exploitation and exploration (Barack et al., 2017; Pearson, Hayden, Raghavachari, & Platt, 2009; Pearson, Heilbronner, Barack, Hayden, & Platt, 2011). In the context of creativity, Kounios et al. (2006) reported increased activity in the PCC during the period leading up to an insightful solution. PCC activity during this period might reflect processes than mediate the shift from exploration (i.e., finding alternative solutions) to exploitation (i.e., focus on an insightful solution). Given that the PCC is involved in detecting changes in the environment and mediating subsequent changes in behavior (Pearson et al., 2011), it may be that the PCC helps detect changes in the overall value of ideas during creative cognition (see Barack et al., 2017), and mediates shifts between different brain networks.

Importantly, the insula, ACC, OFC, and locus coeruleus nucleus are highly interconnected, suggesting that transitions between phasic and tonic LC-NE modes could be associated with activity in the salience network (ACC and insula). The OFC and ACC send major cortical inputs to the locus coeruleus (Aston-Jones & Cohen, 2005a; Porrino & Goldman-Rakic, 1982); the OFC projects to the insula (Aston-Jones & Cohen, 2005a), which also projects to the OFC and ACC (Aston-Jones & Cohen, 2005a-b). These neuroanatomical interconnections raise the possibility that value computations drive LC-NE activity, which, in turn, mediates interactions and transitions between various brain networks. That is, the diffuse projections of the LC-NE system through cortical regions might play a central role in governing network dynamics (Guedj, Meunier, Meunier, & Hadj-Bouziane, 2017), and this idea is consistent with the view that the fMRI BOLD signal may reflect neuromodulatory effects more than changes in the spiking rate of neurons (Logothetis, 2008; Toussay, Basu, Lacoste, & Hamel, 2013).

The idea that LC-NE activity might drive network dynamics is also consistent with other models of LC-NE function. LC-NE activity has been proposed to facilitate network resetting, such that when the LC-NE system is activated, it resets the system by interrupting existing functional networks and facilitating the emergence of new ones (Bouret & Sara, 2005; Guedj et al., 2017; Sara, 2009; see also Mittner et al., 2016). For example, it may be that norepinephrine released by the locus coeruleus resets the attention networks to promote adaptive shifts in attention and changes in behavioral responses (Corbetta, Patel, & Shulman, 2008; Sara & Bouret, 2012). During creative cognition, such attention resetting might facilitate the transition from exploration to exploitation. Integrating these theories of LC-NE function is beyond the scope of the current paper, but with the present framework we hope to stimulate future work that bridges LC-NE function, creative cognition, and value-based decision making.

## Limitations and future directions

By synthesizing ideas and findings from multiple fields, the present framework offers a novel account of creative cognition. However, several issues remain to be addressed. First, this framework assumes that creative cognition is not qualitatively different from normal cognition, in that decision processes that underlie everyday choices are assumed to also support creative processes. However, creative and normal cognition could be seen as mutually exclusive, partially overlapping, or undifferentiated (Abraham, 2013). Clearly, the present framework is incompatible with the mutual exclusivity account, but future work should explore whether the processes underlying creative cognition and economic choice are partially or completely overlapping. Second, in its current conceptualization, this framework does not distinguish between the various aspects or types of creativity (e.g., divergent thinking, insight problem solving, combination of remote semantic associations; see also constrained versus unconstrained cognitive flexibility, Alexander et al., 2007; Hecht et al., 2014). It assumes that the same value-based decision-making processes underlie creative cognition involving all the aforementioned tasks, but future work is required to test this assumption. Third, the current framework has the potential to provide an integrative framework that explains not only creative processes within an individual, but also individual differences in creativity. Clearly, more work is needed to test this aspect of the model. Fourth, this paper has discussed creative generation and evaluation as though these two processes occur largely independently. However, like LC-NE phasic and tonic activity which falls on a continuum, generative and evaluative processes might also fall on a continuum. In addition, it could be that the transitions between these processes might occur too rapidly to be measured using tools that have relatively low temporal resolution (e.g., fMRI). Thus, other neuroimaging methods with greater temporal resolution might be better suited to test some of the framework’s predictions—including the distinctiveness of generation and evaluation as stages in the creative process.

## Conclusion

Recently, several frameworks have been proposed to account for the neural mechanisms that underlie creativity (Boot et al., 2017; Dietrich & Haider, 2016). Unlike previous accounts, our framework draws heavily on neuroeconomics to describe how creative cognition occurs in the brain. By thinking about creative cognition as an adaptive value-maximization process supported by activity in the LC-NE neuromodulatory system, it re-conceptualizes the way we think about the creative process and provides a novel perspective for reinterpreting and integrating existing findings. It also offers many new hypotheses, making the framework highly testable and falsifiable. Although here we have outlined only the key hypotheses, many additional nuanced predictions can be derived from our framework. We believe that this framework can significantly improve our understanding of not just creative cognition, but also the relationship between decision making, neuromodulation, and large-scale brain network dynamics.

